# Phosphatases control the duration and range of cAMP/PKA microdomains

**DOI:** 10.1101/2024.06.04.597457

**Authors:** Filippo Conca, Doruk Kaan Bayburtlu, Mauro Vismara, Nicoletta C. Surdo, Alessandra Tavoni, Leonardo Nogara, Adamo Sarra, Stefano Ciciliot, Giulietta Di Benedetto, Liliana F. Iannucci, Konstantinos Lefkimmiatis

**Affiliations:** Department of Molecular Medicine, University of Pavia, Pavia, Italy; Veneto Institute of Molecular Medicine, 35129 Padova, Italy; Neuroscience institute, Italian National Research Council, Padova, Italy; Department of Biomedical Sciences, University of Padua, Padua, Italy; Department of Pharmaceutical and Pharmacological Sciences, University of Padua, Padua, Italy

## Abstract

The spatiotemporal interplay between the second messenger cyclic AMP (cAMP) and its main effector, Protein Kinase A (PKA) is crucial for the pleotropic nature of this cascade. To maintain a high degree of specificity, the cAMP/PKA axis is organized into functional multiprotein complexes, called microdomains, precisely distributed within the cell. While the subcellular allocation of PKA is guaranteed by a family of tethers called A-Kinase-anchoring Proteins (AKAPs), the mechanisms underlying the efficient confinement of a microdomain’s functional effects are not fully understood. Here we used FRET-based sensors to detect cAMP levels and PKA-dependent phosphorylation within specific subcellular compartments and found that, while free cAMP is responsible for the activation of local PKA enzymes, the dephosphorylating actions of phosphatases dictate the duration of the microdomain’s effects. To test the range of action of PKA microdomains we used rigid aminoacidic nanorulers to distance our FRET sensors from their original location for 10 or 30 nm. Interestingly, we established that cAMP levels do not affect the spatial range of the microdomain while on the contrary, phosphatase activity acts as the main functional boundary for phosphorylated PKA targets. Our findings contribute to the design of a picture where two microdomain-forming events have distinct roles. Cyclic AMP elevations trigger the initial activation of subcellular PKA moieties, while the temporal and spatial extent of the PKA’s actions is regulated by phosphatases.

## Introduction

Protein kinase A (PKA) is a tetrameric enzyme composed by two catalytic (PKACs) and two regulatory (PKARs) subunits. In humans, PKARs are encoded by four genes (RIα, RIβ, RIIα, and RIIβ), while PKACs can be found in three isoforms (Cα, Cβ, or Cγ)(*1*). PKA holoenzymes that contain RI subunits are defined as type I, while type II are the ones that contain RII regulatory subunits. In response to rises in intracellular cAMP levels, the regulatory subunits release the PKAC monomers, which are constitutively active and able to phosphorylate numerous endogenous proteins. Despite the apparent linearity of the signalling pathway and the promiscuous nature of the catalytic subunits, the cAMP/PKA axis exhibits remarkable functional specificity of action. Indeed, PKA can pair extracellular and/or intracellular stimuli to distinct cellular responses, even though cAMP production is the common denominator of all these cues. The precise modalities through which stimuli that increment cellular cAMP levels are coupled to specific outcomes are not fully understood. The leading idea suggests that the cAMP pathway is compartmentalized into functional units called microdomains. Thanks to this organization, changes in cAMP levels triggered by specific stimuli impinge on distinct microdomains generating a sort of spatiotemporal code that allows the cells to interpret and precisely respond to the initial stimulus(*2*, *3*).

If distilled to their essence, microdomains could be characterized mainly by two parameters, i.e. their subcellular localization, which determines the molecular targets within their reach, and their activation/inactivation kinetics, determining the duration of their actions. When it comes to cAMP/PKA microdomains, subcellular distribution is determined by a family of tethers, the A-kinase anchoring proteins (AKAPs)(*4*, *5*). AKAPs associate specifically with the regulatory subunits of PKA (preferably RIIs) thanks to a docking and dimerization (D/D) domain located at their N-terminus(*6*, *7*). The cellular domains where AKAPs constrain PKA holoenzymes are defined by targeting sequences able to localize them and, consequently, their associated proteins to specific regions within cells. Importantly, the roles of AKAPs extend beyond the relocation of PKA. In fact, these proteins can aid the formation of complexes with other microdomain constituents such as phosphatases (PPs), adenylyl cyclases (ACs), and phosphodiesterases (PDEs)(*8*, *9*). The coordination of these components allows for efficient management of cAMP and potentially PKA activity within microdomains. For instance, ACs can offer a local cAMP source, while PDEs, with the aid of cAMP buffers(*10*), are thought to be the enzymes that “shape” intracellular cAMP levels. Managing cAMP availability is considered the principal regulatory mechanism of PKA activity. Nevertheless, plenty experimental evidence suggests the involvement of other events in determining the spatiotemporal entity of cAMP/PKA microdomains(*11*–*13*). It is expected that in response to cAMP rises and PKA activation, the targets within the reach of the microdomain will be phosphorylated. However, lowering the levels of the messenger will result in decreased kinase activity but is not directly linked to the dephosphorylation of the PKA targets. In fact, the definite termination of the functional outcomes of the cascade requires phosphatases, another regulator of microdomains, that dephosphorylate the PKA substrates and effectively end its actions. In line with these considerations, we recently found that PKA-dependent phosphorylation persists longer within domains with low phosphatase activity and subsides earlier where phosphatases are more active(*11*, *13*). Based on these observations, it could be reasonable to assume that the range of action of a kinase-driven microdomain may depend on the phosphatase activity of its surroundings. This is particularly relevant for cAMP/PKA microdomains, as AKAPs bind only the PKARs while PKACs are released in response to cAMP elevations. In principle, free PKACs can diffuse and phosphorylate targets along their path, however this would go against the principle of compartmentalization, unless the diffusion and/or activity of the kinase were controlled. Three modalities that could limit the diffusion of PKACs have been proposed. The first is based on observations suggesting that, at least for AKAP79, even in the presence of cAMP, anchored PKA holoenzymes remain intact and PKACs do not diffuse away from the anchoring sites(*14*). According to this hypothesis, low cAMP elevations fail to induce the release of PKACs, whilst allowing the phosphorylation of proximal targets. Subsequent work used a different approach and proposed an alternative mechanism of free PKAC restriction that is based on the excess of regulatory subunits when compared to the catalytic ones(*15*). Taking in consideration that the regulatory subunits can exhibit two states, one with high affinity for cAMP, and another with high affinity for the catalytic subunits, the abundance of PKARs as compared to the PKACs (up to 17-fold depending on the cell type(*15*, *16*)) could indeed support high rates of association of PKA holoenzymes and limit their diffusion. However, the efficient restriction of PKACs would also depend on the state of PKARs and the levels of cAMP within the microdomain. In fact, when the cAMP levels are high (e.g., during the activation of a microdomain), PKA holoenzymes remain dissociated leaving open the question of how the actions of PKACs are delimited. Finally, N-terminal myristoylation was found to relocate free PKAC by facilitating their membrane association. This modality however increases, rather than decrease, PKA activity near membranes at least in neurons(*12*).

We reasoned that another viable mechanism for restricting the range of PKACs actions, and that of their phosphorylated targets, could rely on phosphatases. In previous work, we demonstrated that PKA-dependent phosphorylation persists longer near the membranes through a mechanism involving the accessibility of phosphatases(*11*, *13*). Based on these findings, we hypothesized that, as PKACs diffuse away from a membrane microdomain, the exposure of their phosphorylated targets to phosphatases will increase, facilitating dephosphorylation and consequently limiting the range of action of the cAMP/PKA domain. To test this hypothesis, we compared two cellular models that we found display radically different cAMP handling modalities, the cervical cancer cell line (HeLa) and the osteosarcoma cell line (U-2 OS). We used rigid peptide nanorulers of known length (10 and 30nm)(*17*) to distance from their original sites the mitochondrial and plasma membrane (PM) versions of the AKAR4(*18*) FRET-based sensor, that monitor PKA-dependent phosphorylation. We found that the vicinity to membranes directly associated with more pronounced and persistent PKA-dependent phosphorylation of the sensors and therefore longer lasting local effects. Interestingly, when the distance of the sensors was increased, the effects of phosphatases became more marked, and the rate of dephosphorylation increased independently of the levels of cAMP. Our data suggest that PKACs and their substrates, once released from membrane bound AKAPs, encounter environments with increasing phosphatase pressure that limit their actions and therefore define both the duration and range of action of PKA microdomains.

## Results

### HeLa and U-2 OS cells display distinct cAMP hydrolyzation kinetics

The intracellular levels of cAMP are defined by a complex multiparametric equilibrium established between intracellular buffers (regulatory subunits and effectors), producers (adenylyl cyclases (ACs)) and the hydrolases responsible for its degradation (PDEs)(*10*). The main intracellular buffers are the free PKA regulatory subunits, which are more abundant(*15*) and display significantly higher affinity than the holoenzymes(*10*, *19*, *20*). While the role of cAMP buffers in the maintenance of low intracellular cAMP levels at basal conditions may be relevant, their high affinity suggests that during cAMP-generating events buffers will be fast saturated, allowing the holoenzymes to be activated. Based on these considerations, whether a physiological signal will reach the cAMP concentration threshold to activate PKA mainly depends on its ability to tilt the balance in favor of production(*21*). To study the involvement of PDE activity in cAMP/PKA compartmentalization, we employed the cAMP-sensitive FRET-based sensor EPAC-S^H187^ (H187)(*22*, *23*). We performed a preliminary screening on different cell lines and chose to continue our investigation using HeLa and U-2 OS cells based on their wide use in the field(*11*, *22*) and differences in cAMP handling. As shown in **fig 1A**, we designed an experimental protocol aiming to test the balance between cAMP production and degradation thanks to the use of different dosages and combinations of AC activators and PDE inhibitors. HeLa cells expressing H187 responded very blandly to low doses of the AC activator forskolin (FSK) while the addition of 8-Methoxymethyl-3-isobutyl-1-methylxanthine (IBMX) to block PDEs produced striking responses, suggesting that cAMP hydrolysis due to PDEs is very pronounced in these cells (**fig 1B**). Increasing the levels of FSK (20µM) in the presence of IBMX resulted in the saturation of the sensor. We used the same protocol in U-2 OS cells and, as shown in **fig 1C**, low doses of FSK alone resulted in a higher FRET change, while PDE inhibition (IBMX) had only a bland effect of the FRET signal that was further enhanced by higher FSK doses. These experiments suggested that PDEs are crucial regulators of cAMP signals in HeLa cells, while are less active in U-2 OS. To directly test the cAMP hydrolyzing potential of HeLa and U-2 OS cells, we used an experimental protocol where high doses of FSK in the presence of IBMX were used to induce the saturation of the sensor, followed by rinsing of IBMX (whilst keeping the levels of FSK constant) to release PDE activity but maintain the high production levels (**fig 1D**). As shown in **fig 1E & F**, treatment with high doses of FSK and IBMX produced fast, saturating responses in both cell lines. On the other hand, rinsing IBMX, to release PDE activity, resulted in drastically different kinetics in the two models. In line with our previous results, in HeLa cells the FRET signal reversed with very fast kinetics, despite the persistent FSK-dependent activation of ACs (maximal cAMP production), confirming high PDE activity in these cells. On the contrary, the same maneuver (IBMX removal) resulted in a minimal response in U-2 OS cells, indicating that the release of PDE activity was not enough to overcome the AC-dependent cAMP production. In fact, in U-2 OS cells the FRET signal recovered only when FSK was rinsed away from the experimental medium (**fig 1F**).

The H187 FRET sensor bares a Q270E point mutation in the EPAC module which increases the affinity of the sensor for cAMP approximately 2.5-fold, making this construct particularly fit for measuring low levels of cAMP(*21*, *22*). High-affinity sensors constitute also potential buffers and are less optimal for measuring variations in cells that can retain higher basal cAMP levels. To exclude an effect of H187 overexpression, we repeated the experiments shown in **fig 1A &D**, using a different FRET-based sensor that lacks the Q270E substitution called EPAC-S^H84^ (H84)(*24*) and therefore displays lower affinity for cAMP. As shown in **Supplementary fig 1A & B**, albeit an overall lower dynamic range, H84 displayed kinetics that virtually overlap with the responses measured by H187 in both, HeLa and U-2 OS. Taken together these experiments indicate that in HeLa cells PDE activity is strongly impeaching on cytosolic cAMP levels contrary to U-2 OS, where the PDE-dependent hydrolysis is less dominant and cAMP production by ACs is scarcely contrasted.

### Different mechanisms control intracellular cAMP levels in HeLa and U-2 OS

In our experiments, cAMP production was initiated at the plasma membrane, by FSK, a labdane diterpene that activates specifically all transmembrane ACs, except AC9(*25*) and the soluble AC10 (sAC)(*26*). Under these conditions, it is expected that cAMP will be higher at the PM and its free levels will likely decrease due to buffering and PDE activity as it diffuses towards cytosolic sites. The main intracellular cAMP buffers are the free PKA regulatory subunits(*17*, *27*). As shown in **Supplementary fig 2A**, in HeLa cells the expression of PKA regulatory subunits type I (RI) is more pronounced as compared to U-2 OS cells, while less evident are the differences in the expression of RII subunits. These data, together with the striking difference in PDE activity (**fig 1**), would suggest that the cAMP signal “shaping” capacity will be more pronounced in HeLa than U-2 OS. To test whether the dynamics of cAMP signalling at the proximity of PM are different than those observed in the cytosol we repeated the experimental protocols depicted in **fig 1A** using sensors targeted to the PM. Due to its high affinity, the sensor H187 was deemed less optimal for measurements at the PM where cAMP levels are expected to be high. We employed a PM-targeted version of the FRET-based sensor EPAC-S^H30^ (H30) that displays similar kinetics and sensitivity to H84 **Supplementary fig 2B & C**. Using cells expressing PM-EPAC-S^H30^ (PM-H30), we were unable to observe different kinetics near the plasma membrane than those measured in the cytosol in any of the two cell models. More specifically, in HeLa cells, low levels of FSK (5µM) resulted in marginal cAMP elevations (virtually overlapping with the cytosolic responses of the soluble sensors H30, H84 and H187), while PDE inhibition with high doses of IBMX (500µM) strongly affected the FRET signal, suggesting that PDEs impinge on cAMP production on site (**fig 2A**). On the other hand, in U-2 OS expressing PM-H30, treatment with FSK 5µM, triggered a significant FRET change while the effects of PDE inhibition remain limited, as also observed in the cytosol (**fig 2B**). To estimate the PDE activity in the vicinity of the PM, we used the protocol of **fig 1D**, in HeLa and U-2 OS expressing PM-H30, and FRET changes specifically due to cAMP hydrolysis were registered. As shown in **fig 2C & D**, IBMX removal resulted in fast extended responses in HeLa cells but had nearly no effect in U-2 OS, which signals reversed only when FSK was removed from the experimental medium.

**Figure 1.**
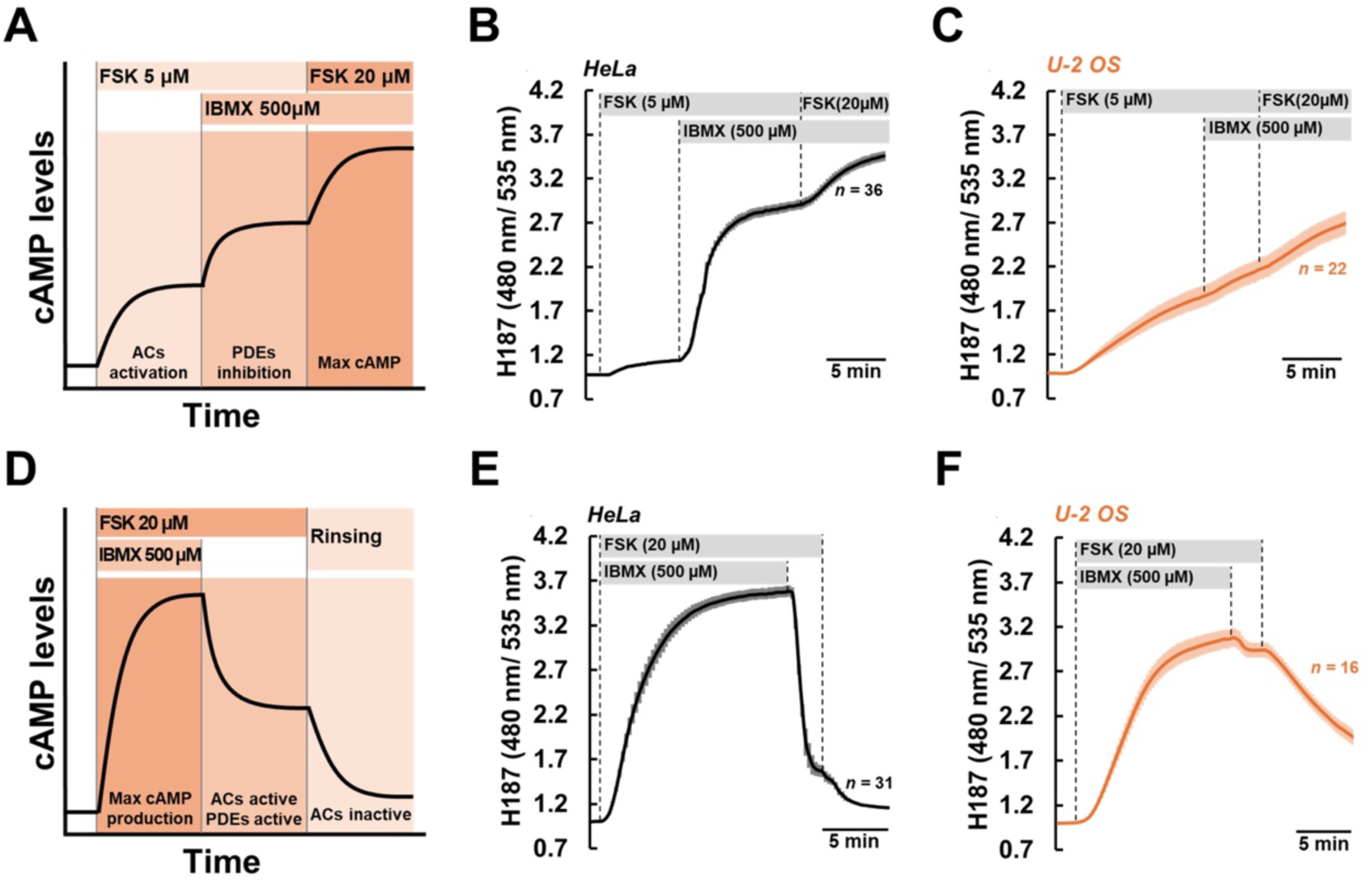
Hela and U-2 OS exhibit different PDE-dependent cAMP hydrolysis potential. **(A)** Schematic representation of the experimental strategy used in (B & C). Hela cells **(B)** or U-2 OS cells **(C)** expressing the cAMP-sensitive FRET-based sensor H187 were challenged with low doses (5µM) of forskolin (FSK) alone or combined with IBMX (500µM) to inhibit PDEs, followed by addition of high doses of FSK (20µM) to achieve saturation of the sensor. Contrary to U-2 OS, HeLa cells did not respond to low FSK unless PDEs were blocked, suggesting that these cells harbour high cAMP-hydrolyzing activity (traces are expressed as the average ± SEM; n=36 Hela cells; n=22 U-2 OS). **(D)** Schematic representation of the experimental strategy used in (E & F). Hela cells **(E)** or U-2 OS cells **(F)** were challenged with FSK (20µM) combined to IBMX (500µM), resulting in the saturation of the sensor. Subsequent rinsing of IBMX, to unmask the IBMX-sensitive PDE activity resulted in the recovery of the FRET signal. In line with (B &C) in Hela cells FRET recovery, which mirrors the rate of cAMP hydrolysis, was significantly faster, suggesting higher PDE activity as compared to U-2 OS (traces are expressed as the average ± SEM; n=31 Hela cells; n=16 U2-OS).

**Figure 2.**
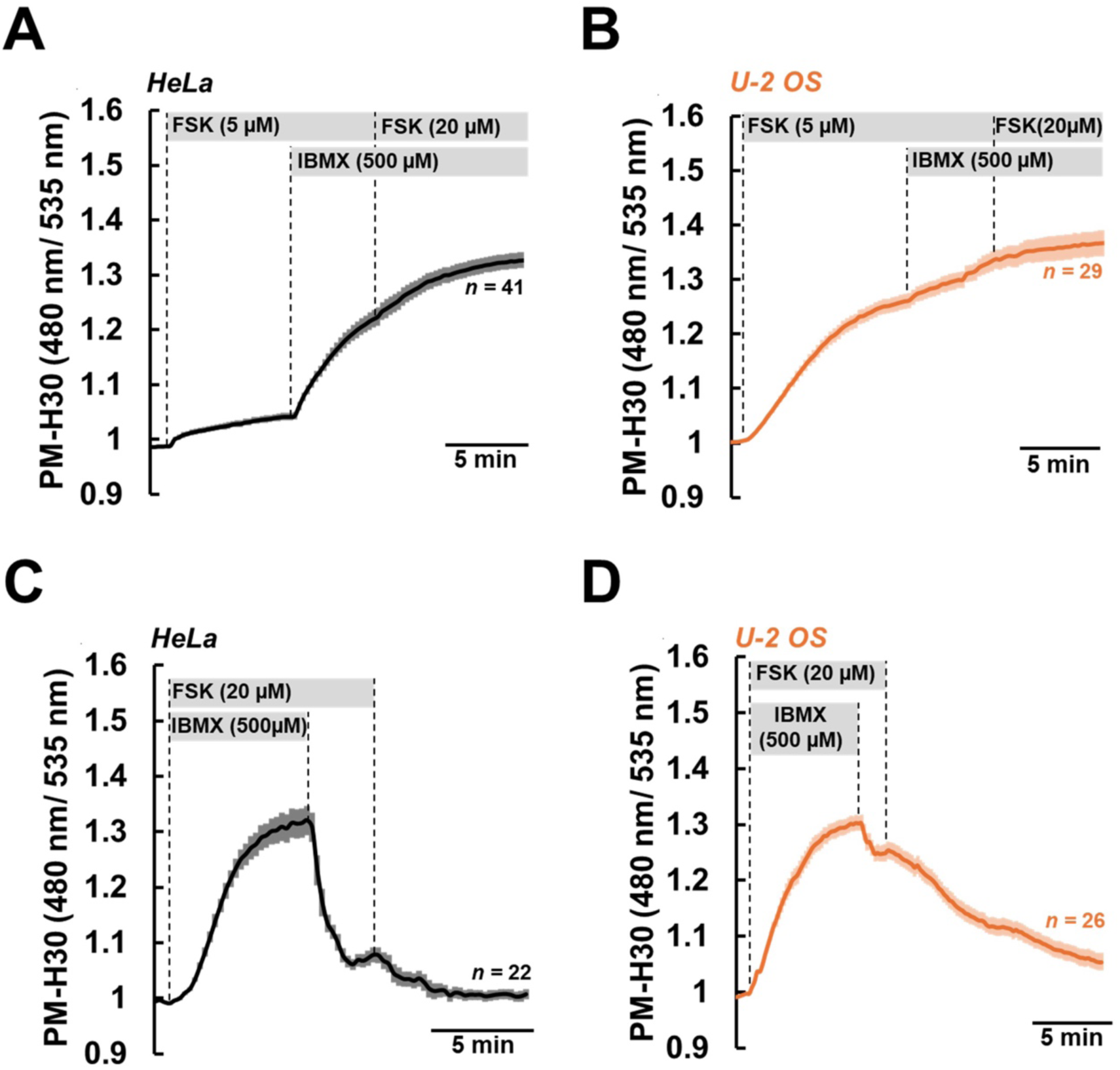
In Hela but not U-2 OS PDE activity shapes the sub-plasma membrane cAMP levels. Hela cells **(A)** or U-2 OS cells **(B)** expressing a plasma membrane targeted version of the cAMP-sensitive FRET-based sensor EPAC-S^H30^ (PM-H30) were challenged with low doses (5µM) of forskolin (FSK) alone or combined with IBMX (500µM) to inhibit PDEs, followed by addition of high doses of FSK (20µM) to achieve saturation of the sensor. In U-2 OS, inhibition of PDEs had only marginal effects, on the contrary in HeLa cells addition of IBMX resulted in a significant FRET response (traces are expressed as the average ± SEM; n=41 Hela cells; n=29 U2-OS). Hela **(C)** or U-2 OS cells **(D)** expressing PM-H30 were treated with saturating concentrations of FSK (20µM) and IBMX (500µM). Rinsing of IBMX to release PDE activity had no effect in U-2 OS cells, but resulted in a fast decrease of the FRET signal in HeLa cells (traces are expressed as the average ± SEM; n=22 Hela cells; n=26 U2-OS).

Taken together, these experiments suggested that PDEs differentially impinge on both, the kinetics and the levels of free cAMP at the plasma membrane. In fact, in cells with high (HeLa) or low (U-2 OS) PDE activity, AC activation using low levels of FSK produced drastically different cAMP responses. Moreover, when we compared cAMP dynamics in different compartments of HeLa and U-2 OS cells, we found that cAMP kinetics in the cytosol mirrored those measured at the plasma membrane in both cell models. Interestingly, in our experiments we did not observe any cAMP gradients independently of the cAMP hydrolyzing ability of each cell line. Our data are in line with recent reports suggesting that PDE activity alone is not sufficient for the generation of intracellular cAMP gradients(*28*, *29*).

### PKA activation in specific compartments mirrors intracellular cAMP levels

Among the cAMP effectors, PKA is the one that most relies on compartmentalization. In fact, tethering of this kinase to specific sub-cellular locations by AKAPs is at the basis of the generation of cAMP microdomains. Both the affinity for cAMP and AKAPs of the PKA holoenzymes depend on their regulatory subunits. Type II regulatory subunits display lower cAMP affinity but higher tendency to complex with AKAPs, while RIs are more sensitive to the messenger but less prone to complex with the tethers(*30*). In addition, recent experimental evidence, suggests that type I PKARs may act as cAMP buffers and could contribute to its compartmentalization(*27*, *31*). Based on these considerations, we checked the expression levels of the cAMP/PKA signalling components in HeLa and U-2 OS. Western Blotting experiments showed that both cell models express RI and RII subunits as well as the main catalytic subunit (PKACα) and the main phosphatases (**Supplementary fig 2A**). To determine whether the different cAMP kinetics observed in HeLa and U-2 OS cells resulted in the activation of distinct cAMP/PKA domains, we measured the kinetics of PKA activation in different subcellular locations using targeted versions of the FRET-based sensor AKAR4(*18*), which provides a good approximation of the PKA-dependent substrate phosphorylation dynamics. We opted for two different compartments, the mitochondria and plasma membrane, and as control we used the soluble AKAR4 parent sensor. The outer mitochondrial membrane was chosen because it hosts a cAMP/PKA microdomain built around D-AKAP2(*32*) and subject to specific regulatory modalities(*11*). Moreover, it was recently proposed that PKA microdomains build around D-AKAP2 may be important for cAMP buffering and diffusion(*10*). On the other hand, the plasma membrane compartment was chosen because it hosts most of the ACs and therefore is expected to be the domain with the highest cAMP levels. We used two well calibrated sensors, OMM-AKAR4(*11*) targeted by a generic tag to the outer mitochondria membrane, and the lipid raft-binding Lyn-AKAR4 as a PM-targeted sensor(*18*).

To test how cAMP production and hydrolysis affects PKA-dependent phosphorylation in different compartments, we subjected HeLa and U-2 OS cells to the experimental protocol depicted in **fig 3A**. As shown in **fig 3B**, the soluble AKAR4 sensor responded differently in HeLa and U-2 OS cells, faithfully mirroring the cAMP kinetics previously measured by H187 in the two cell lines (**fig 1B&C**). Similarly, treatment with low doses of FSK (5µM) resulted in increased PKA-dependent phosphorylation at the outer mitochondrial membrane only in U-2 OS cells (**fig 3C**). Finally, addition of IBMX to the experimental medium resulted in strong (saturating) activation of both AKAR4 and OMM-AKAR4 (**fig 3B&C**). These results confirm that PKA activation in the cytosol and at the outer mitochondria membrane depends on the cAMP levels, and are in line with previous reports indicating that these compartments do not present differences in their cAMP managing ability(*11*, *13*). Contrary to the soluble and OMM sensors, Lyn-AKAR4 responded to FSK 5µM in both HeLa and U-2 OS cells (**fig 3D**). Interestingly, in U-2 OS, saturation of Lyn-AKAR4 was achieved independently of PDE inhibition, while in HeLa cells addition of IBMX induced a further response of Lyn-AKAR4, confirming the superior hydrolyzing ability of these cells. These experiments confirmed the importance of cAMP managing for the activation of compartmentalized PKA. Indeed, in HeLa cells, where cAMP is kept under strict control by PDEs, addition of IBMX was necessary for releasing PKA-dependent phosphorylation induced by low doses of FSK. Interestingly, this was evident in all compartments, except the plasma membrane, where the penetrance of PDE activity was partial, probably due to the vicinity to the cAMP producing sites. On the other hand, in U-2 OS cells, where PDE activity was negligible, all treatments resulted in saturating PKA responses further confirming the importance of cAMP hydrolysis in PKA activation.

**Figure 3.**
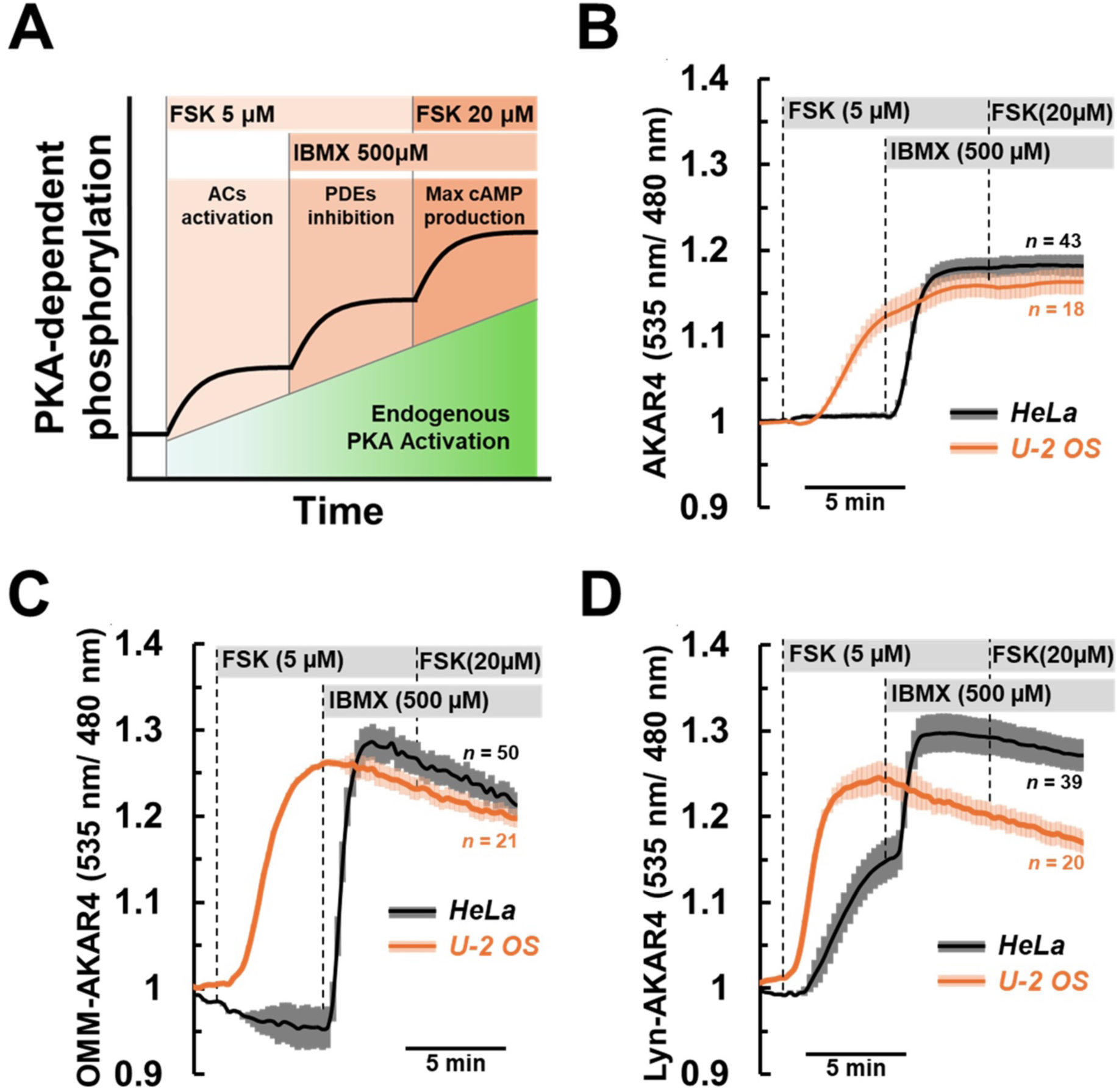
PKA-dependent phosphorylation of different domains is dictated by global cAMP levels in Hela and U-2 OS cells. **(A)** Schematic representation of the experimental strategy used. **(B)** Hela cells (dark traces) and U2-OS cells (light traces) expressing the PKA-dependent phosphorylation FRET-based sensor AKAR4, its mitochondrial targeted version OMM-AKAR4 **(C)**, or its Lipid raft targeted version Lyn-AKAR4 **(D)**, were treated with low doses (5µM) of Forskolin (FSK) alone or combined with IBMX (500µM), followed by addition of FSK (20µM) to saturate the sensor. PKA-dependent phosphorylation kinetics mirrored those of the cAMP levels, except for the plasma membrane domain in HeLa cells (traces are expressed as the average ± SEM; AKAR4: n=43 Hela cells and n=18 U2-OS; OMM-AKAR4: n=21 Hela cells and n=50 U2-OS; Lyn-AKAR4: n=39 Hela cells and n=20 U2-OS).

### PKA termination dynamics are similar in HeLa and U-2 OS cells

PKA microdomains are defined as subcellular locations where the kinase is tethered by AKAPs in the vicinity of both its targets and the machinery that controls its activation kinetics and range of action(*33*). Our experiments demonstrated that cAMP increases drive PKA activation, which is then translated to target phosphorylation and cellular responses. However, the inverse does not apply, since PKA deactivation in response to cAMP hydrolysis, limits the ex novo phosphorylation of targets but does not affect those already phosphorylated. In fact, to terminate the cellular responses induced by PKA is necessary to dephosphorylate its targets through the actions of phosphatases(*11*, *13*, *34*). Based on these considerations, it could be suggested that PKA substrate dephosphorylation is a complex process, initiated by the inactivation of the kinase, but completed by the actions of the phosphatases, It is therefore the rate of dephosphorylation that defines the extent of cellular responses induced by cAMP/PKA microdomains.

To test the importance of phosphatases on the termination of local PKA-dependent phosphorylation we employed the experimental protocol shown in **fig 4A**. HeLa and U-2 OS cells expressing different versions of the AKAR4 sensor were challenged with high doses of FSK and IBMX, a treatment that leads to the saturation all sensors. We then supplemented the experimental medium with H-89, an ATP analog that is well known to inhibit PKA and was expected to tilt the balance to the favor of phosphatases. As shown in **fig 4B**, AKAR4 responded to treatment with FSK (20µM) and IBMX (500µM) with a fast increase in FRET that reached saturation within 2-3 minutes. Addition of H-89 (60µM) that strongly inhibited endogenous PKA catalytic subunits resulted in the dephosphorylation of the sensor, which was extremely fast and virtually overlapping in the two cell lines. We then repeated this protocol in cells expressing OMM-AKAR4. As shown in **Fig 4C**, the dephosphorylation kinetics observed in both cell lines were drastically slower than the ones measured by the soluble sensor, and once more, were very similar in U-2 OS and HeLa cells. Interestingly, the kinetics of dephosphorylation observed at the lipid rafts under the plasma membrane using Lyn-AKAR4 in HeLa and U-2OS cells were slightly different and fell in between the rates measured for AKAR4 and OMM-AKAR4 (**fig 4D**), nevertheless displaying no significant difference between the two cell models. Taken together these experiments suggest that the activation kinetics of PKA faithfully mirror changes in cAMP levels. On the other hand, the dephosphorylation of PKA substrates both targeted and soluble, which determines the duration of the kinase’s actions, depends on phosphatase activity, and is decoupled from the messenger levels, in fact, the termination kinetics were overlapping in both HeLa and U-2 OS despite their different cAMP managing modalities.

**Figure 4.**
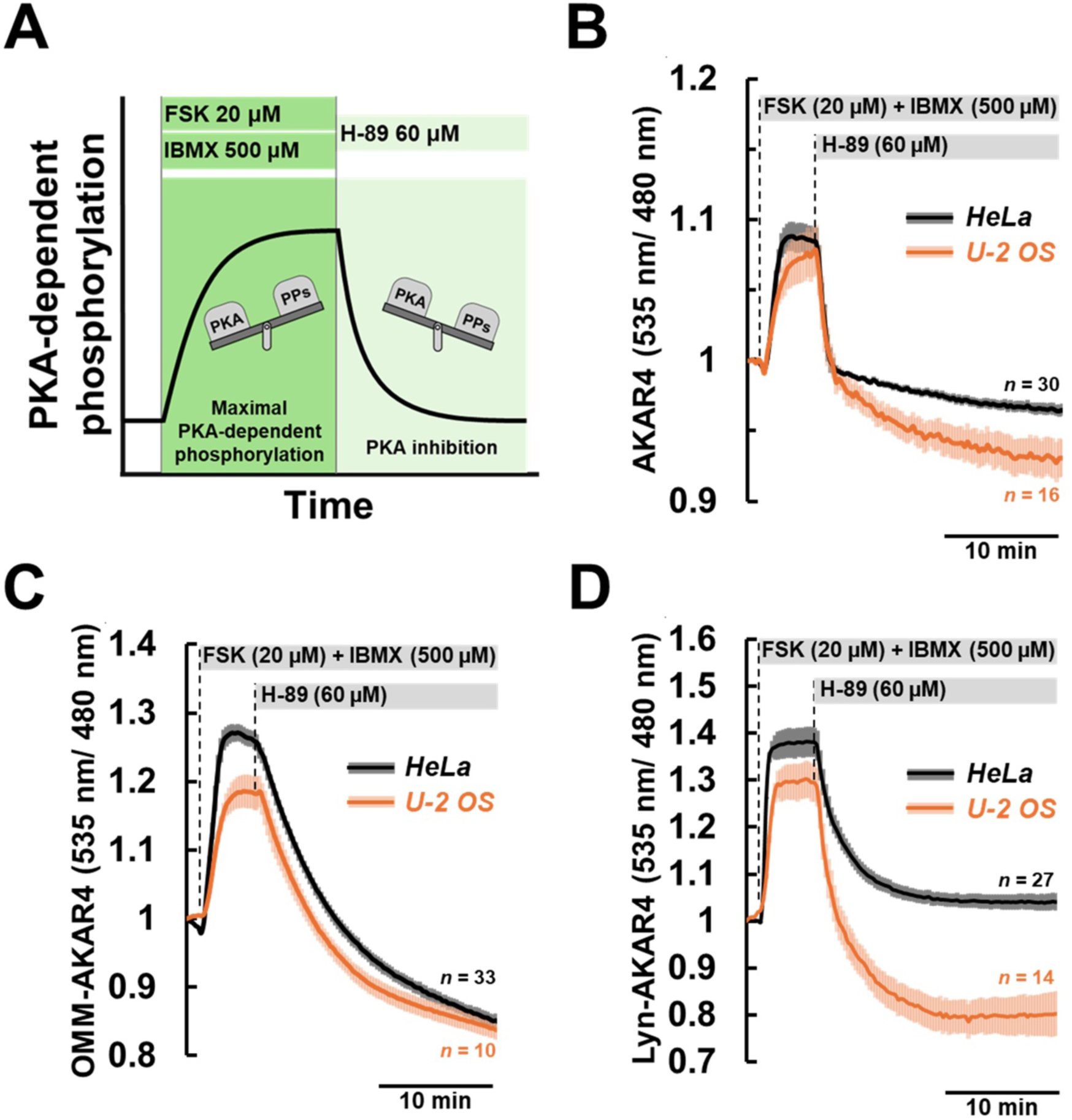
Hela and U-2 OS cells have similar PKA substrate dephosphorylation kinetics. **(A)** Schematic representation of the experimental strategy used. **(B)** Hela cells (dark traces) and U2-OS cells (light traces) expressing AKAR4, OMM-AKAR4 **(C)**, or Lyn-AKAR4 **(D)**. Cells were treated with high concentrations of FSK (20µM) and IBMX (500µM) until all sensors reached saturation. At this point the PKA inhibitor H-89 (60µM) was also added to the experimental medium. The rate of dephosphorylation within different compartments was virtually overlapping independently of the cell model (traces are expressed as the average ± SEM; AKAR4: n=30 Hela cells and n=16 U2-OS; OMM-AKAR4: n=33 Hela cells and n=10 U-2 OS; Lyn-AKAR4: n=27 Hela cells and n=14 U2-OS).

### Phosphatases can define the range of OMM-related PKA-dependent events

Our previous experiments demonstrated that different compartments could display various degrees of phosphatase activity. This is in line with work from our group in primary neonatal cardiac myocytes, showing that PKA substrates at the OMM are less accessible to phosphatases and therefore, stay phosphorylated for longer periods as compared to cytosolic targets (*11*). These data point to the phosphatases as the main regulators of the duration of PKA-dependent events.

The OMM is the interface of mitochondria with the cytosol and its resident proteins, many of which are PKA substrates, participate to important processes including mitochondrial dynamics and mitophagy(*35*, *36*). Interestingly, our data may suggest that proteins reaching the mitochondria will encounter decreasing phosphatase pressure, while for those leaving the organelles the phosphatase pressure will increase. To test this possibility and establish the range within which phosphatases could influence the distribution of phosphorylated PKA substrates, we designed a series of AKAR4-based sensors exploiting rigid spacers of known length(*17*). As shown in **Supplementary fig 3A**, we sandwiched the SAH10 (10nm) and SAH30 (30nm) peptides between the targeting sequence and the sensor. This way we could control the distance of the active AKAR4 core from the surface of mitochondria. As expected, adding SAH10 and SAH30 had no effect on the targeting of OMM-AKAR4 and both OMM-SAH10-AKAR4 and OMM-SAH30-AKAR4 localized strictly at the mitochondria **Supplementary fig 3Bi**. All versions of the sensor retained their ability to measure PKA-dependent phosphorylation and dephosphorylation kinetics, as demonstrated in experiments using HeLa cells co-expressing OMM-AKAR4, OMM-SAH10-AKAR4 or OMM-SAH30-AKAR4 together with an mCherry-tagged PKA catalytic subunit. As shown in **Supplementary fig 3C**, control cells expressing just mCherry, responded to FSK/IBMX treatment reaching saturation (**panel i**), while H-89 treatment evidence low basal, unsolicited PKA activity to all constructs except Lyn-SAH10-AKAR4 and Lyn-SAH30-AKAR4 (**panel ii**). When PKAC-mCherry was expressed, cells did not respond to FSK/IBMX, likely due to saturation (**panel iii**), while H-89 treatment at the basal level generated large responses (**panel iv**), demonstrating their sensitivity to endogenous phosphatases. Finally, as summarized in **supplementary fig 3D**, all sensors responded with FRET-ratio changes to PKAC overexpression, suggesting that the characteristics of the parent sensor were not altered by the addition of spacers.

Next, we used these sensors to estimate the reach of OMM-based cAMP/PKA microdomains in cells with high (HeLa) and low (U-2 OS) PDE activity. To determine the eventual contribution of ACs, PDEs and phosphatases, we designed a protocol based on sequential maneuvers of AC activation, PDE inhibition and PKA inhibition (**supplementary fig 3E**). As expected, in HeLa cells (**fig 5A**), activation of ACs using high doses of forskolin produced only marginal responses that were greatly increased in after PDE inhibition using IBMX, independently of the sensor version. Once the sensors reached their saturation, we added H-89 to the experimental medium that shifted the balance in favor of the phosphatases and resulted in the recovery of the FRET signal. This maneuver kept cAMP levels high but inhibited the catalytic PKA-subunits. Interestingly, the recovery kinetics of different sensors were not the same, with soluble AKAR4 being the fastest, the OMM-AKAR4 being the slowest while OMM-SAH10-AKAR4 and OMM-SAH30-AKAR4 were somewhat in between. To quantify the differences observed, we calculated the tau-parameter which is the time constant of the exponential decay of a curve for each sensor. As summarized in **fig 5B,** we found that the dephosphorylation kinetics of OMM-SAH10-AKAR4 and OMM-SAH30-AKAR4 became faster when the distance from the mitochondria increased. We next employed the same protocol in U-2 OS cells expressing the different versions of the AKAR4-based sensors. As shown in **fig 5C**, high doses of FSK suffice to saturate all the sensors, independently of PDE-inhibition, while H-89 addition induced the recovery of the FRET signals with kinetics that were the fastest in the soluble fraction (AKAR4) the slowest at the mitochondria (OMM-AKAR4), while became progressively faster by increasing the distance from the mitochondria (**fig 5D**). All together, these experiments link the activity of phosphatases to the range of PKA-dependent phosphorylation signals and suggest that dynamic dephosphorylation of PKA substrates can define both the duration and range of PKA-dependent microdomains at the outer mitochondrial membrane.

**Figure 5.**
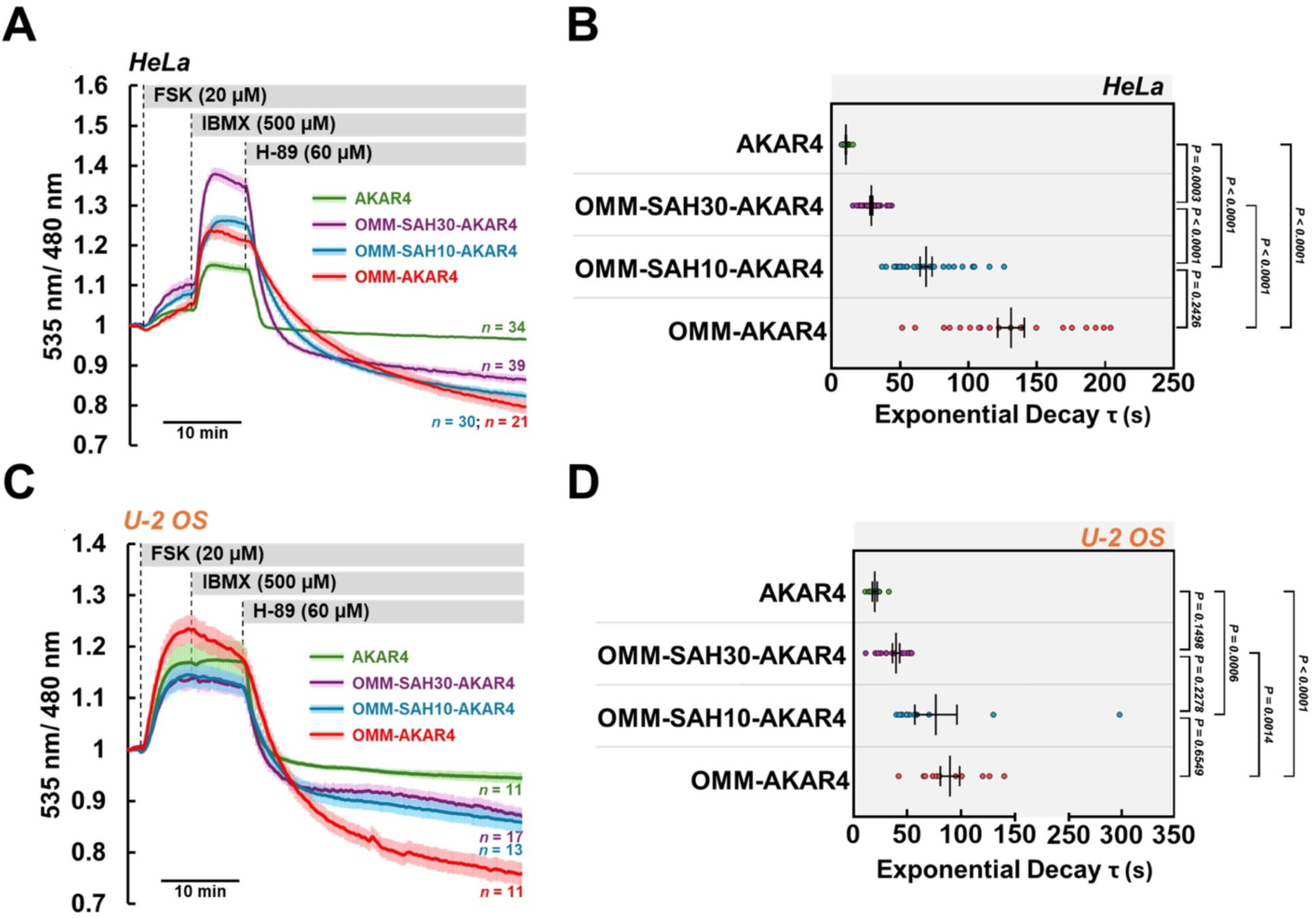
Phosphatase-dependent dephosphorylation of PKA substrates increases proportionally to the distance from mitochondria. **(A)** HeLa cells expressing AKAR4 (green trace), OMM-SAH30-AKAR4 (violet trace), OMM-SAH10-AKAR4 (blue trace) or OMM-AKAR4 (red trace). All sensors were saturated by treatment with FSK (20µM) combined to IBMX (500µM). Addition of H-89 (60µM) in the presence of FSK/IBMX, resulted in different dephosphorylation kinetics as measured by the recovery of the FRET signals. **(B)** The exponential decay of each sensor curve was assessed using Tau (data expressed as the average ± SEM; AKAR4: n=34; OMM-AKAR4: n=39; OMM-SAH10-AKAR4: n=30; OMM-SAH30-AKAR4: n=21). Similar experiments were performed in U2-OS cells **(C)** and Tau descriptor of the exponential decay of the curves is shown in **(D)** (data expressed as the average ± SEM; AKAR4: n=11; OMM-AKAR4: n=11; OMM-SAH10-AKAR4: n=13; OMM-SAH30-AKAR4: n=17). In both cellular models, Tau decreases were inversely proportional to the distance from the outer mitochondrial membrane, indicating increasing phosphatase pressure moving away from the organelles. P values indicated, Seconds (s).

### Phosphatases impinge on the range of sub-plasma membrane PKA-dependent events

Contrary to the outer mitochondrial membrane where cAMP kinetics faithfully mirror those in the cytosol(*11*), domains in the vicinity of plasma membrane contain higher cAMP levels (**fig 3D**) that could allow PM-PKA moieties to respond with different kinetics. This possibility prompt us to test the effects of phosphatases on the range of PM-PKA-dependent events. We generated and validated modified versions of the Lyn-AKAR4 containing the SAH10 and SAH30 spacers (Lyn-SAH10-AKAR4, Lyn-SAH30-AKAR4) (**Supplementary fig 3Bii** and validated in **Supplementary fig 3C&D**). As shown in **fig 6A**, in HeLa cells Lyn-AKAR4 responded strongly to high levels of FSK and reached saturation in response to IBMX. Interestingly, the responses of Lyn-SAH10-AKAR4 and Lyn-SAH30-AKAR4 to FSK were not as marked as those of their parent sensor, while IBMX addition resulted in a more evident increase in PKA-dependent phosphorylation, suggesting a stronger involvement of PDEs. Inhibition of PKACs by H-89 caused the recovery of the FRET-signal in all sensors. As summarized in **fig 6B**, the distance from the plasma membrane associated to the rate of dephosphorylation of the sensors (tau value). However, the differences were evident to a lesser extent from those observed at the OMM. Specifically, adding SAH10 or SAH30 resulted in a faster dephosphorylation rate when compared to Lyn-AKAR4, however the differences between these two constructs were not significant (**fig 6B**). Similar results were obtained also in U-2 OS cells as shown in **fig 6C** and summarized in **fig 6D**.

**Figure 6.**
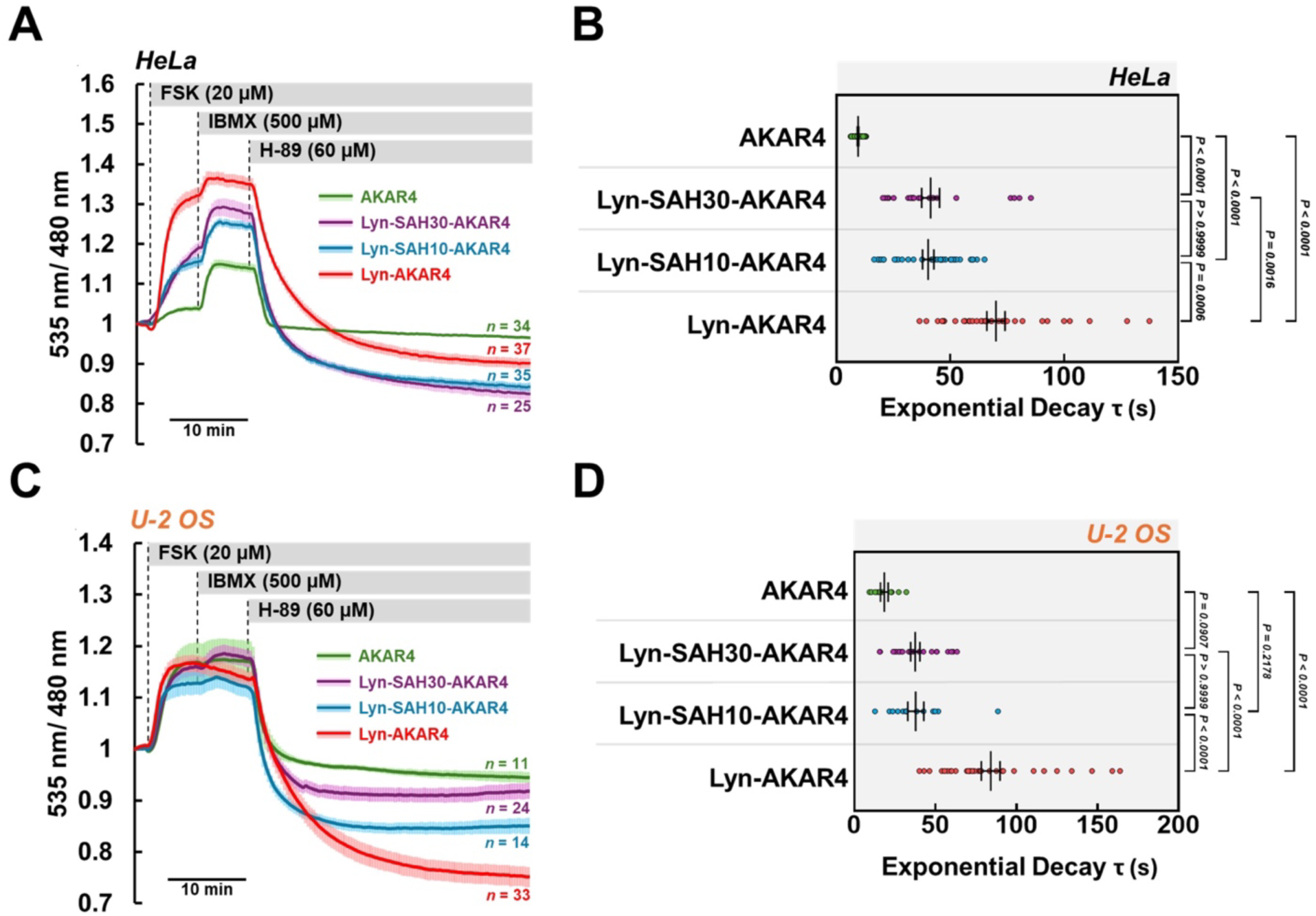
Phosphatase-dependent dephosphorylation of PKA substrates is affected by the distance from the plasma membrane. **(A)** HeLa cells expressing AKAR4 (green trace), Lyn-SAH30-AKAR4 (violet trace), Lyn-SAH10-AKAR4 (blue trace) or Lyn-AKAR4 (red trace). All sensors were saturated by treatment with FSK (20µM) combined to IBMX (500µM). Addition of H-89 (60µM) in the presence of FSK/IBMX, resulted in different dephosphorylation kinetics as measured by the recovery of the FRET signals. **(B)** The exponential decay of each sensor curve was assessed using Tau (data expressed as the average ± SEM; AKAR4: n=34; Lyn-AKAR4: n=37; Lyn-SAH10-AKAR4: n=35; Lyn-SAH30-AKAR4: n=25). Similar experiments were performed in U2-OS cells **(C)** and Tau descriptor of the exponential decay of the curves is shown in **(D)** (data expressed as the average ± SEM; AKAR4: n=11; Lyn-AKAR4: n=33; Lyn-SAH10-AKAR4: n=14; Lyn-SAH30-AKAR4: n=24). In both cellular models, Tau indicated that increases in phosphatase activity reached a plateau already after 10nm from the plasma membrane. P values indicated, Seconds (s).

## Discussion

PKA compartmentalization is a complex task that requires the coordination of two types of machineries, one that guarantees the vicinity of the kinase to its designated targets and another in charge of its activation/deactivation kinetics. Cyclic AMP is generally produced at the plasma membrane and must reach intracellular PKA moieties by diffusion. The levels of cAMP able to reach a specific microdomain depend on multiple factors, notably PDEs(*17*) and intracellular buffers(*10*). PKARs type I are the most relevant cAMP buffers as they are produced in excess (∼17fold) compared to the PKACs(*15*, *16*) and when free display both, very high affinity for cAMP(*21*) and may redistribute to generate biomolecular condensates that act as buffers(*27*).

In the general notion of PKA microdomains, the structural basis of PKA segregation within specific compartments represents a paradox. In fact, while AKAPs offer a structural platform for the tethering of PKA tetramers, they do so by complexing with the regulatory subunits of the tetramer. This feature, however, does not ensure the strict confinement of the PKACs that, in response to cAMP elevations, are released from the regulatory subunits and become free to diffuse away from their site of activation. Recently, it was suggested that AKAP-bound PKA tetramers remain intact independently of their activation status(*14*). However, this model cannot be generalized, as it was tested specifically for AKAP79-based complexes. Moreover, this model and the general idea that PKACs remain linked to PKARs when cAMP levels are high, were challenged by subsequent studies(*12*, *15*). Several mechanisms that limit PKAC diffusion have been proposed, including myristoylation(*12*) and binding to excess regulatory subunits(*15*, *31*). Despite these mechanisms, it is unequivocal that, once released from the tetramer, the degree of freedom of PKACs increases and, consequently, the probability to encounter and phosphorylate substrates that are not strictly part of the specific microdomain. In this manuscript, we investigated the existence of mechanisms that can limit the range of action of PKA microdomains by acting on the free PKACs and their phosphorylated substrates. We reasoned that the regulation of cAMP/PKA microdomains goes beyond their activation/inactivation kinetics and set to understand the distinct contributions of cAMP levels and phosphatases in determining the temporal and spatial range of a microdomain’s actions.

We used two cellular models, HeLa and U-2 OS, on the basis of their dramatically different management of the messenger. HeLa cells displayed high PDE activity that alone was sufficient to regulate the activation of virtually all cAMP microdomains measured, including those near the plasma membrane. On the contrary, U-2 OS relied exclusively on the production of cAMP by ACs, with PDEs being of marginal regulatory importance. Our experiments did not evidence any cAMP gradients and we found that the messenger kinetics measured at the production site were faithfully mirrored in the cytosol, independently of the cell type and therefore regulatory mechanism. We concluded that, in the two cell models used, generalized cAMP increases/decreases were the sole determinants of PKA activation/inactivation in all the subcellular compartments investigated. Using soluble and targeted versions of AKAR4, a FRET-based sensor that mirrors the phosphorylation/dephosphorylation kinetics of PKA substrates(*11*), we demonstrated that the termination pace of PKA-dependent events was determined by the accessibility of phosphatases within the microdomain. In addition, measurements of PKA-dependent phosphorylation persistence at known distances from specific domains (mitochondria and PM) suggested that phosphatase pressure increases in direct relationship to the distance from the microdomain. Our data, together with recently published results(*11*, *17*, *37*) support a model where the balance between cAMP production, buffering and, most importantly, PDE-dependent hydrolyzation, is responsible for the activation and deactivation kinetics of localized PKA moieties. Specifically, based on the levels of PDEs and buffers positioned in its vicinity, each microdomain will be activated in response to different cAMP elevations. Similarly, the higher the hydrolyzing capacity of a location, the more efficiently cAMP levels will drop and PKA will return to its inactive tetrameric state. Translating of cAMP-induced PKA activation to cell function requires, and is driven by, the reversible phosphorylation of substrates. Interestingly, once a target is phosphorylated, its regulation is decoupled from the levels of the messenger and the status of the kinase. In fact, cellular responses triggered by PKA activation will subside only after dephosphorylation of the substrates, thanks to the actions of phosphatases. We suggest that the rate of target dephosphorylation depends on the spatial distribution of phosphatases and likely determine both the temporal and spatial range of a microdomain’s actions. Based on our data, PKACs released from the outer mitochondrial membrane, or the plasma membrane (as well as possible phosphorylated targets) will encounter environments with increasing phosphatase pressure that contribute to the confinement of the microdomain’s reach.

## Materials and methods

### Reagents

Forskolin (FSK, Cat. No. 11018), 3-isobutyl-1-methylxantine (IBMX, Cat. No. 13347) and H-89 dihydrochloride (H-89, Cat. No. 10010556) were purchased from Cayman Chemical (USA). Dimethyl sulfoxide (DMSO, Cat No. D8418), phosphate-buffered saline (PBS, Cat. No. P4417), Tween 20 (Cat. No. P1379), bovine serum albumin (BSA, Cat. No. A9647), and skim milk powder (Cat. No. 70166) were purchased from Merck (Germany).

### Cell culture and transfection

U-2 OS and HeLa cells were grown at 37 °C in a 5% CO_2_ atmosphere in Dulbecco’s modified Eagle’s medium with High Glucose (DMEM, Cat. No. ECM0728, Euroclone, Italy) supplemented with 10% fetal bovine serum (Cat. No. ECS5000L, Euroclone, Italy), 100 U/ml penicillin, 100 μg/ml streptomycin (Cat. No. ECB3001D, Euroclone, Italy), and 2 mM L-Glutamine (Cat. No. ECB3000S, Euroclone, Italy). Cells were split every 2–3 days at a confluence of 80–90%. Twenty-four hours before transfection, cells were plated onto custom 15 mm diameter circular glass coverslips (thickness 0.16 mm, purchased from Vemi S.r.l., Italy) and were allowed to grow to 50% of confluence. Cells were transfected with Lipofectamine 2000 Reagent (Cat. No. 11668019, Thermo Fisher Scientific, USA) following manufacturer’s instructions.

### Plasmids and cloning strategies

The construct mCherry2-C1 was a gift from Michael Davidson (Addgene plasmid # 54563; http://n2t.net/addgene:54563; RRID:Addgene_54563), pcDNA3-AKAR4 was a gift from Jin Zhang (Addgene plasmid # 61619; http://n2t.net/addgene:61619; RRID:Addgene_61619) and pcDNA3-Lyn-AKAR4 was a gift from Jin Zhang (Addgene plasmid # 61620; http://n2t.net/addgene:61620; RRID:Addgene_61620) all above constructs were purchased from Addgene. The *pcDNA3-OMM-AKAR4* sensor and pcDNA3-PKA-mCherry were previously generated in our laboratory(*11*). The *pcDNA3-OMM-SAH10-AKAR4* and *pcDNA3-OMM-SAH30-AKAR4* constructs were obtained by PCR amplification of *SAH10* and *SAH30* respectively and then cloning in *pcDNA3-OMM-AKAR4* using NheI and BamHI restriction sites. *SAH10* and *SAH30* were amplified from *pcDNA3-Epac1-camps-SAH10-PDE4A1* and *pcDNA3-Epac1-camps-SAH30-PDE2Cat* kindly gifted by Professor Martin J. Lohse, Dr. Andreas Bock and Dr. Isabella Maiellaro. The *pcDNA3-SAH10-AKAR4* and *pcDNA3-SAH30-AKAR4* were obtained from *pcDNA3-OMM-SAH10-AKAR4* and *pcDNA3-OMM-SAH10-AKAR4* respectively, by excision of the OMM targeting sequence using the double HindIII restriction site. The constructs *pcDNA3-Lyn-SAH10-AKAR4* and *pcDNA3-Lyn-SAH30-AKAR4* were obtained from *pcDNA3-OMM-SAH10-AKAR4* and *pcDNA3-OMM-SAH30-AKAR4* respectively. The OMM targeting sequence was excised by HindIII digestion. Primers containing the Lyn sequence and the HindIII restriction site were then used as insert in the ligation reaction.

### FRET-based imaging

U-2 OS and HeLa cells plated onto 15 mm diameter circular glass coverslips (thickness 0.16 mm, purchased from Vemi S.r.l., Italy) and transfected with different FRET sensors were mounted onto an open perfusion chamber RC-25F (Warner Instruments, USA). The cells were bathed in Ringer’s modified buffer: 125 mM NaCl; 5 mM KCl; 1 mM Na_3_PO_4_; 1 mM MgSO_4_; 20 mM Hepes; 5.5 mM D-(+)-glucose; 1 mM CaCl_2_; pH adjusted to 7.4 using 1M NaOH. The experiments were performed on an Olympus IX83 inverted microscope (Olympus, Japan) equipped with Filter Wheel mode with Optospin (Cairn Research Ltd, England) and a CCD camera Retiga R^TM^ Series R1 (Teledyne Photometrics, USA). The cyan fluorescent proteins (Cerulean for AKAR4 or mTurquoise for H187) were excited for 200 milliseconds at 430 nm, while the emission fluorescence was collected every 15 s for both donor and acceptor fluorophores at 480 and 535 nm, respectively. Automatic image collection and pre-liminary analysis were performed using the MetaFluor® 7.10.5.476 (Molecular Devices, USA). Raw data were transferred to Excel (Microsoft, USA) for background subtraction and generation of the ratios. Graphs were generated with Excel (Microsoft, USA), or GraphPad Prism8 software (GraphPad, USA).

### Western blotting

The cells were lysed in cold Lysis buffer supplemented with cOmplete mini EDTA-free protease inhibitor cocktail (Cat. No. 11836170001, Roche, Switzerland) and phosphatases inhibitor PhosSTOP™ (Cat. No. 04906837001, Roche, Switzerland). The cell lysates were cleared at 12.000 rpm at 4 °C for 10 min and 40 μg of proteins were loaded onto 10% polyacrylamide gel (Tris-glycine gels, Thermo Fisher Scientific, USA) and then transferred onto polyvinylidenefluoride (PVDF) membranes (Cytiva, UK). Subsequently, the membranes were blocked for 1 h at room temperature with 10% (w/v) non-fat-dry milk in 0,2% TBS-Tween (TBS-T) and then incubated overnight at 4 °C with primary antibody (1:1000) in 5% non-fat-dry milk-TBS-T. The day after, membranes were washed three times with TBST at room temperature and then incubated for 1 h at room temperature with peroxidase-conjugated secondary antibodies. Peroxidase activity was detected with enhanced chemiluminescence (Luminata Crescendo Western HRP, Cat. No. WBLUR0500, Merck, Germany). Membranes were stripped using Restore Western Blot Stripping Buffer (Cat. No. 46430 Thermo Fisher Scientific) for 15 min at room temperature and thoroughly washed with TBS-T.

Anti-PKA Catα (C-20) (sc-903; Santa Cruz Biotechnologies, USA), anti-PKA Reg Iα/β (G-6) (sc-271446; Santa Cruz Biotechnologies, USA), anti-PKA Reg IIβ (L-16) (sc-26803; Santa Cruz Biotechnologies, USA), anti-PP1 (E-9) (sc-7482; Santa Cruz Biotechnologies, USA), anti PP2Aα/β (ab32141; Abcam, UK), and anti β-actin (A3854; Merck, Germany). Anti-Mouse (115-035-003) and Anti-Rabbit (111-035-003) peroxidase-conjugated secondary antibodies were purchased from Jackson ImmunoResearch Europe (UK). Anti-goat peroxidase-conjugated secondary antibody (sc-2033) was purchased from Santa Cruz Biotechnology (USA).

### Data fitting

Datapoints following the addition of FSK and IMBX have been fitted using a lineas function to obtain a slope (dratio/dt). To perform the exponential decay fit, individual traces have been subtracted for the slope of a linear fit performed on the last 50 points to obtain a flat baseline, allowing a reliable single exponential decay fitting using the following equation: Y=(Y0 - Plateau)*exp(-K*X) + Plateau, (initial values equal to 1, 1000 iterations), the time constant tau (t) is obtained. Slopes and exponential decay data have been analyzed using GraphPad Prism (version 10.0.2 for Mac, GraphPad Software, Boston, Massachusetts USA), performing a Saphiro-Wilk normality test followed by the ROUT outliers identification test (Q=1%). The statistical analysis performed is an ordinary One-way ANOVA and a Tukey’s multiple comparison test. Data are expressed as mean±SEM.

### Indirect immunofluorescence and confocal imaging

HeLa cells were seeded onto 15 mm diameter circular glass coverslips (thickness 0.16 mm, purchased from Vemi S.r.l., Italy) and transfected with *pcDNA3-AKAR4, pcDNA3-OMM-SAH30-AKAR4, pcDNA3-OMM-SAH10-AKAR4, pcDNA3-OMM-AKAR4, pcDNA3-Lyn-SAH30-AKAR4, pcDNA3-Lyn-SAH10-AKAR4,* and *pcDNA3-Lyn-AKAR4*. The day after, cells were fixated in 4% paraformaldehyde-PBS (sc-281692, Santa Cruz Biotechnology, USA) for 10 minutes, then washed 5 times in PBS and permeabilized in 0.5% Triton X-100 (Cat. No. X100, Merck, Germany)-TBS for 10 min. During blocking, cells were incubated at room temperature in 0.1% Triton-X100-, 2% BSA-, 0.1% Sodium Azide (Cat. No. 71289, Merck, Germany)-TBS. The primary antibody anti-Citrate Synthase (D7V8B), purchased from Cell Signalling Technology (USA, Cat. No. 14309S) diluted 1:300 in the same blocking solution as well as the secondary antibody Alexa Fluor 647 (Cat. No. A21244, Life Technologies, USA). Plasma membranes were stained with wheat germ agglutinin (WGA) conjugated with Alexa Fluor 647 (Cat. No. W32466, Life Technologies, USA) following manufacturer instructions. Cells immunostained for Citrate Synthase were acquired on a Leica TCS SP8 confocal scanning microscope using oil immersion 63x (HC PL APO 63x/1.40 Oil CS2, Leica, Germany) objective. Cells stained with WGA were acquired with a Zeiss LSM900 Confocal upright confocal scanning microscope (Carl Zeiss Microscopy, Germany) using an oil immersion 63x (Zeiss Plan-Apochromat 63x/1.4, Carl Zeiss Microscopy, Germany) objective.

### Statistics

FRET experiments were independently repeated at least three times with similar results. The sample size of each experiment is reported in the figure legends. The overall FRET graphs are shown as average of single-cell traces ± SEM. Data were analyzed with GraphPad Prism8 software (GraphPad, USA). For confocal fluorescence experiments each coverslip, at least 10 random cells were acquired across 2 or 3 independent experiments. The fluorescence intensity was calculated with Fiji (NIH, USA) and the graphs were generated with GraphPad Prism8 software (GraphPad, USA).

## Acknowledgments

We thank the core facility “Centro Grandi Strumenti” (CGS) at the University of Pavia for providing access to the Confocal Microscopy laboratory in particular Amanda Oldani and Patrizia Vaghi for the technical support. We would like to thank Professor Martin J. Lohse, Dr. Andreas Bock and Dr. Isabella Maiellaro for sharing the SAH10 and SAH30 constructs. We are grateful to funding from Human Frontier Science Program Research Grant (HSFP # RG0024/2022), the AIRC foundation for cancer research (grant # IG2021 ID 26140) and AFM Telethon research grant (#25089) to KL.

## Contributions

Performed experiments FC, DKB, MV, NCS, AT, AS, SC. Analyzed and visualized data FC, DKB, LN. Conceptualization, experimental design FC, DKB, LFI, GDB, KL. Writing of the first draft FC, LI, KL. All authors contributed to manuscript editing.

## Supplementary Figures

**Supplementary figure 1.**
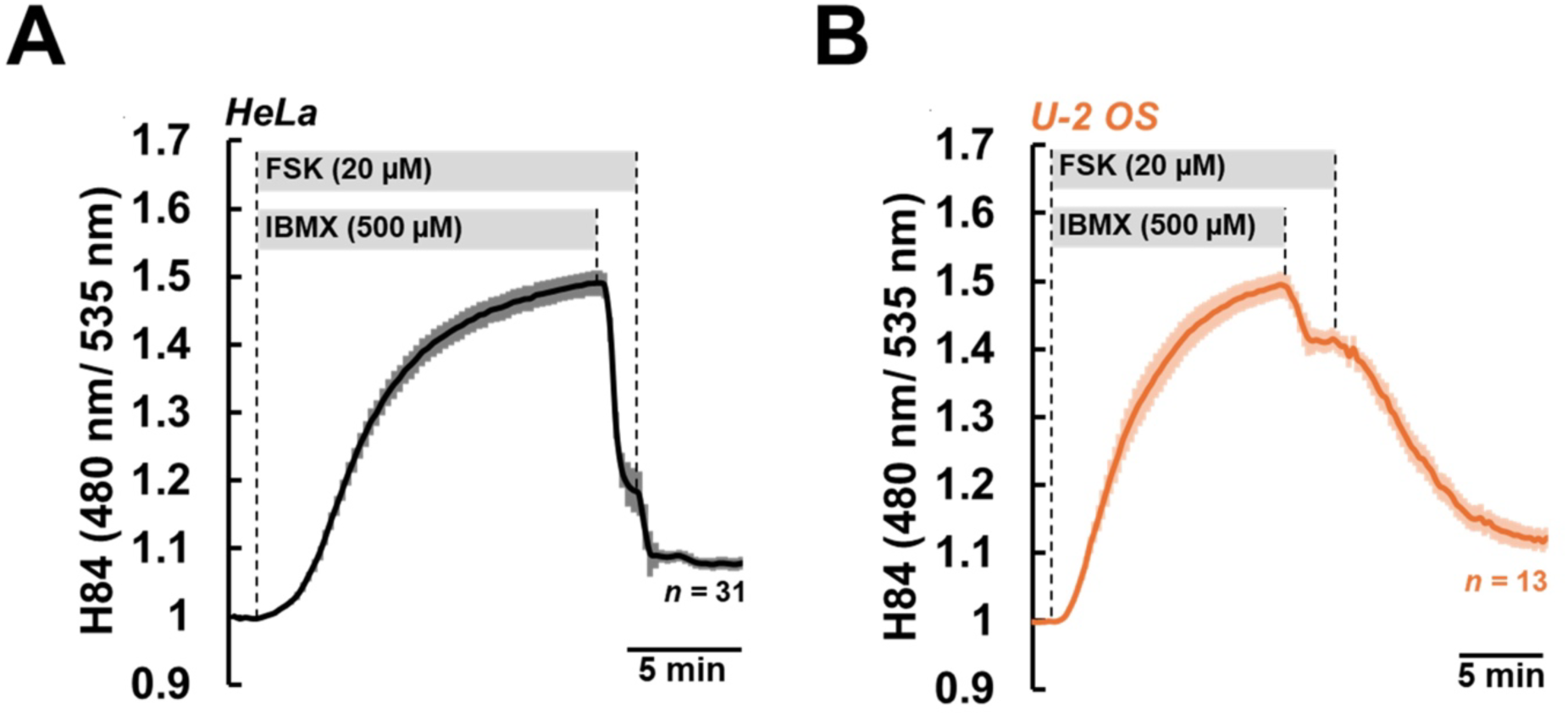
Characterization of the cAMP sensitive FRET-based sensor EPAC-S^H84^ (H84). Hela cells **(A)** and U-2 OS cells **(B)** expressing H84 (which lacks the Q270E substitution and shows lower cAMP affinity) were treated with high levels of FSK (20µM) combined to IBMX (500µM) to reach saturation. Withdrawal of IBMX to release the inhibition of PDEs, resulted in kinetics that mirrored those measured by H187 (which displays higher cAMP affinity (fig E & F). Data are presented as mean value ± SEM; n=31 Hela cells; n=13 U-2 OS.

**Supplementary figure 2.**
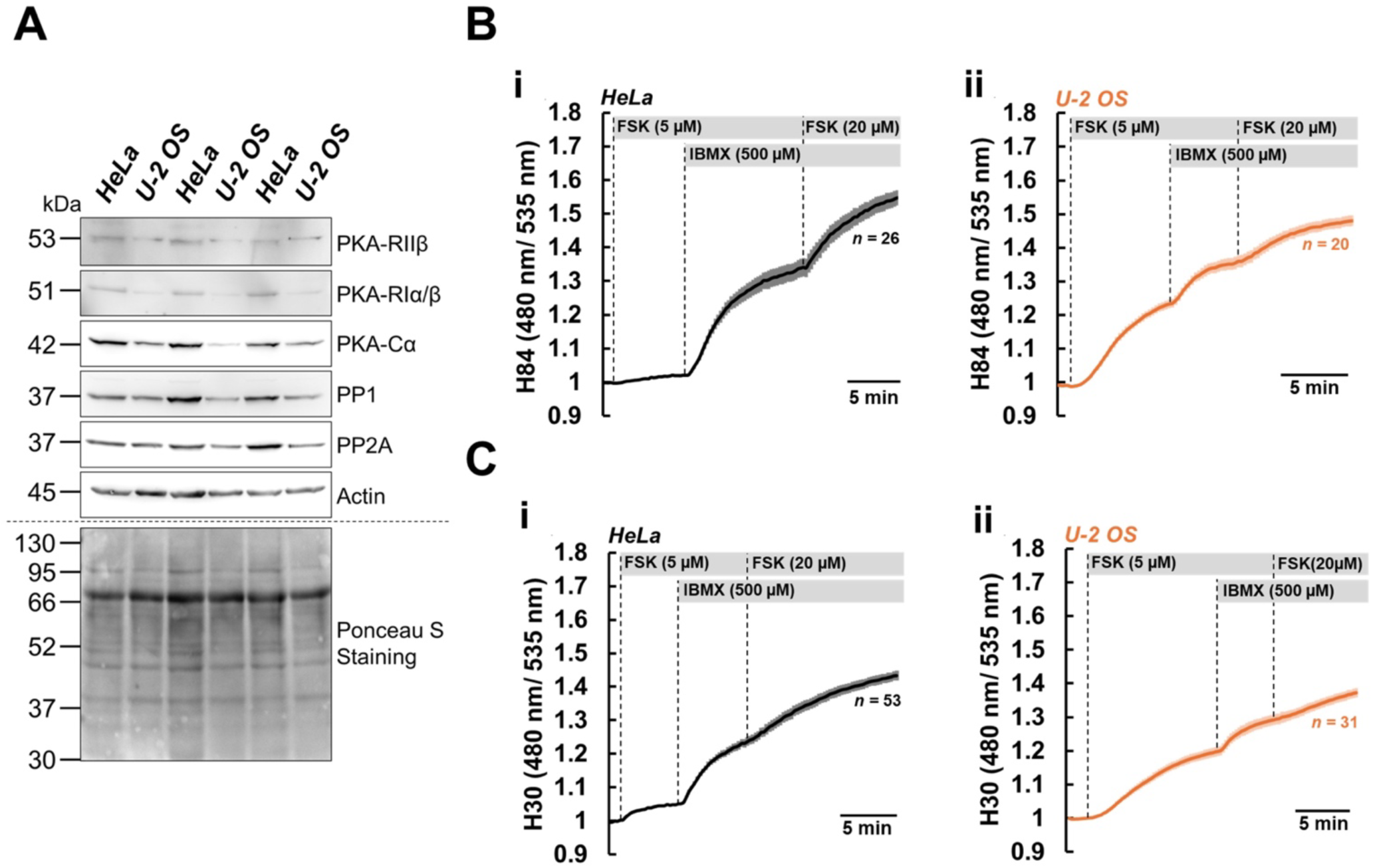
Characterization of cAMP signalling and its components in HeLa and U-2 OS cells. **(A)** Western Blotting using total cell lysates from Hela and U-2 OS. Three biological replicates are presented, and β-actin and Ponceau S staining were used as loading and quality controls. **(B)** Comparison of the FRET-based cAMP sensor EPAC-S^H84^ (H84) in HeLa **(i)** and U-2 OS **(ii)**. **(C)** Comparison of the cAMP sensor EPAC-S^H30^ (H30) in HeLa (i) and U2-OS (ii). Traces are expressed as the average ± SEM; H84: n=26 Hela cells and n=20 U2-OS; H30: n=53 Hela and n=31 U-2 OS.

**Supplementary figure 3:**
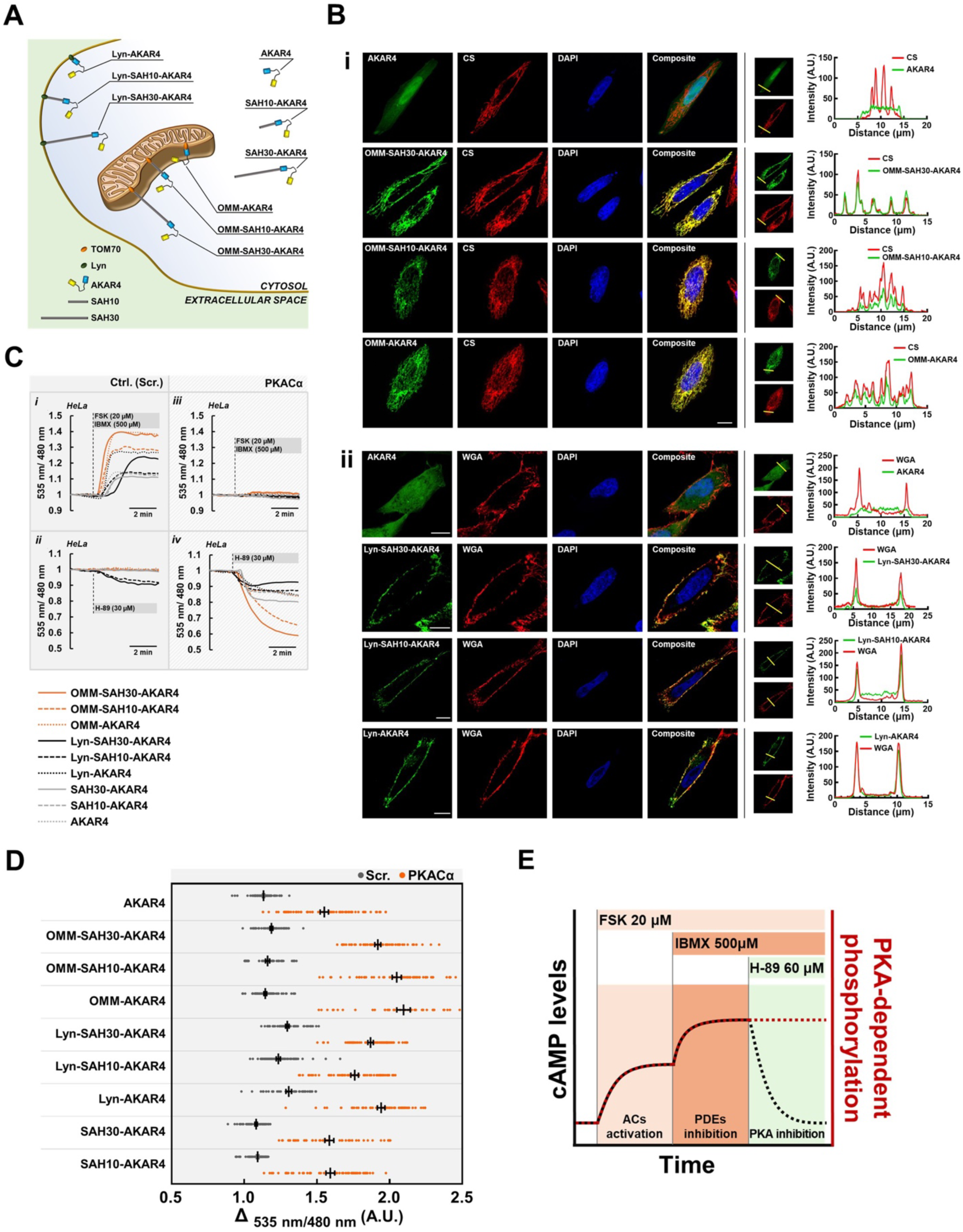
Validation of SAH10 and SAH30-containing OMM-AKAR4 and Lyn-AKAR4 sensors. **(A)** Schematic representation of the different FRET-based sensors developed. **(B)** Confocal photomicrographs of HeLa cells expressing the soluble parent sensor AKAR4, and its outer mitochondrial membrane versions OMM-SAH30-AKAR4, OMM-SAH30-AKAR4, OMM-AKAR4 **(i)** or its plasma membrane targeted versions, Lyn-SAH30-AKAR4, Lyn-SAH10-AKAR4, Lyn-AKAR4 **(ii)**. Insets: line-intensity profiles of the indicated regions. Citrate Synthase (CS) and Wheat Germ Agglutinin (WGA) were used as markers of mitochondria and plasma membrane respectively, while DAPI was used to stain the nuclei (experiments were repeated 4 and 2 times respectively). **(C)** Calibration of all FRET sensors used, cells expressing the indicated constructs alone or together with a catalytically active PKACα and treated with FSK (20µM) together with IBMX (500µM) or alternatively with H-89 (30µM). **(D)** Summary of the basal and maximal FRET ratio for each sensor representing with good approximation the range of each construct. Data are presented as dot plot of individual data points and mean ± SEM error bars; n=97 AKAR4; n=66 AKAR4 + PKACα; n=62 OMM-SAH30-AKAR4; n=59 OMM-SAH30-AKAR4 + PKACα; n=64 OMM-SAH10-AKAR4; n=55 OMM-SAH10-AKAR4 + PKACα; n=67 OMM-AKAR4; n=42 OMM-AKAR4 + PKACα; n=59 Lyn-SAH30-AKAR4; n=62 Lyn-SAH30-AKAR4 + PKACα; n=64 Lyn-SAH10-AKAR4; n=54 Lyn-SAH10-AKAR4 + PKACα; n=43 Lyn-AKAR4; n=57 Lyn-AKAR4 + PKACα; n=62 SAH30-AKAR4; n=51 SAH30-AKAR4 + PKACα; n=58 SAH10-AKAR4; n=55 SAH10-AKAR4 + PKACα; **(E)** Schematic representation of the experimental strategy used for the assessment of the rate of PKA-substrate dephosphorylation in the presence of high cAMP levels.

